# An exploratory study on the microbiome of northern and southern populations of *Ixodes scapularis* ticks predicts changes and unique bacterial interactions

**DOI:** 10.1101/2021.12.17.473158

**Authors:** Deepak Kumar, Latoyia P. Downs, Abdulsalam Adegoke, Erika Machtinger, Kelly Oggenfuss, Richard S. Ostfeld, Monica Embers, Shahid Karim

## Abstract

The black-legged tick (*Ixodes scapularis*) is the primary vector of *Borrelia burgdorferi*, the causative agent of Lyme disease in North America. However, the prevalence of Lyme borreliosis is clustered around the northern states of the United States of America. This study utilized a metagenomic sequencing approach to compare the microbial communities residing within *Ix. scapularis* populations from north and southern geographic locations in the USA. Using a SparCC network construction model, potential interactions between members of the microbial communities from *Borrelia burgdorferi-*infected tissues of unfed and blood-fed ticks were performed. A significant difference in bacterial composition and diversity among northern and southern tick populations was found between northern and southern tick populations. The network analysis predicted a potential antagonistic interaction between endosymbiont *Rickettsia buchneri* and *Borrelia burgdorferi* sensu lato. Network analysis, as expected, predicted significant positive and negative microbial interactions in ticks from these geographic regions, with the genus *Rickettsia, Francisella*, and *Borreliella* playing an essential role in the identified clusters. Interactions between *Rickettsia buchneri* and *Borrelia burgdorferi* sensu lato needs more validation and understanding. Understanding the interplay between the micro-biome and tick-borne pathogens within tick vectors may pave the way for new strategies to prevent tick-borne infections.

## 1. Introduction

Vector-borne diseases affect over one billion people every year and have been expanding alarmingly in recent years [1-3]. Among vector-borne diseases, Lyme disease is one of the most reported infectious diseases in the United States and corresponds to over 90% of vector-borne infections in North America [4-5]. Almost 300,000 cases of Lyme disease are reported every year in the United States, and *Ixodes scapularis* is known as its primary vector. Most Lyme disease cases are clustered in Northeastern and upper Midwest states [6]. *Ix. scapularis* ticks are also prevalent in the Southern states, but the dearth of Lyme disease infections is linked with the restricted distribution and scarcity of the primary *B. burgdorferi* s.l. reservoir, white-footed mice, *Peromyscus leucopus* [7]. *B. burgdorferi* s.l., an infection rate of ∼ 35- 50% have been reported among ticks from the Northeastern states [8-9], whereas tick populations in the southern states are more rarely infected [10-14]. Several studies have reported the microbial composition residing within the ticks [15- 22]. Despite this work, limited information exists on which to base comparisons of the microbiome residing within *Ix. scapularis* from the Lyme disease endemic and non-endemic areas, nor between different tissue types within ticks. Also, there is a dearth of information on the interaction between recently identified tick endosymbionts or commensal microorganisms and *B. burgdorferi* s.l. or pathogenic microbes. Efforts have been made to identify commensal and symbiotic microbes in the tick microbiome [8, 23]. Significant variations associated with geography, species, and sex in *Ixodes* microbiota have been reported [14]. The impact of microbiome communities on physiological processes in ixodid ticks, including reproductive fitness and vector competence, has been suggested [24-29]. In the United States, tick microbiome studies have shown variation in sex, species, and geography. In Canada, *Ix. scapularis* microbiome from eastern and southern regions did not differ significantly concerning the geographic origin, sex, or life stages (Clow et al., 2018). These variable results motivate additional studies to generally detect whether geographic and demographic patterns exist.

This s9tudy conducted a comparative analysis of the microbiome residing within the *Ix. scapularis* populations collected from Lyme disease-endemic and non-endemic areas in the United States utilizing 16S rRNA sequencing. This study also provided an insight into the microbiome composition in unfed and partially-blood-fed tick tissues, including midgut, salivary glands, and ovaries.

## 2. Results

### 2.1 Illumina MiSeq 16S V1-V3 sequencing

The total number of reads obtained for whole tick samples collected from LA, NY, OK, and PA were 2,427,191, corresponding to 2,812 operational taxonomic units (OTUs), a minimum number of reads for a sample was 36,358, while a maximum number of reads for a sample was 114,478. For tick tissues, the total number of reads was 546,024, corresponding to 1,009 OTUs, a maximum number of reads for a sample was 56,785, while the minimum number of reads was 17,258. In our analyses, each tissue sample was rarefied to 5,000 sequences. In comparison, whole tick samples were rarefied to 16,000 sequences. The rarefaction curve (Figure 1) plotted between the number of observed OTUs, and the sequence as mentioned above depth reached plateau suggested sufficient sample coverage for further analysis.

**Figure 1.**
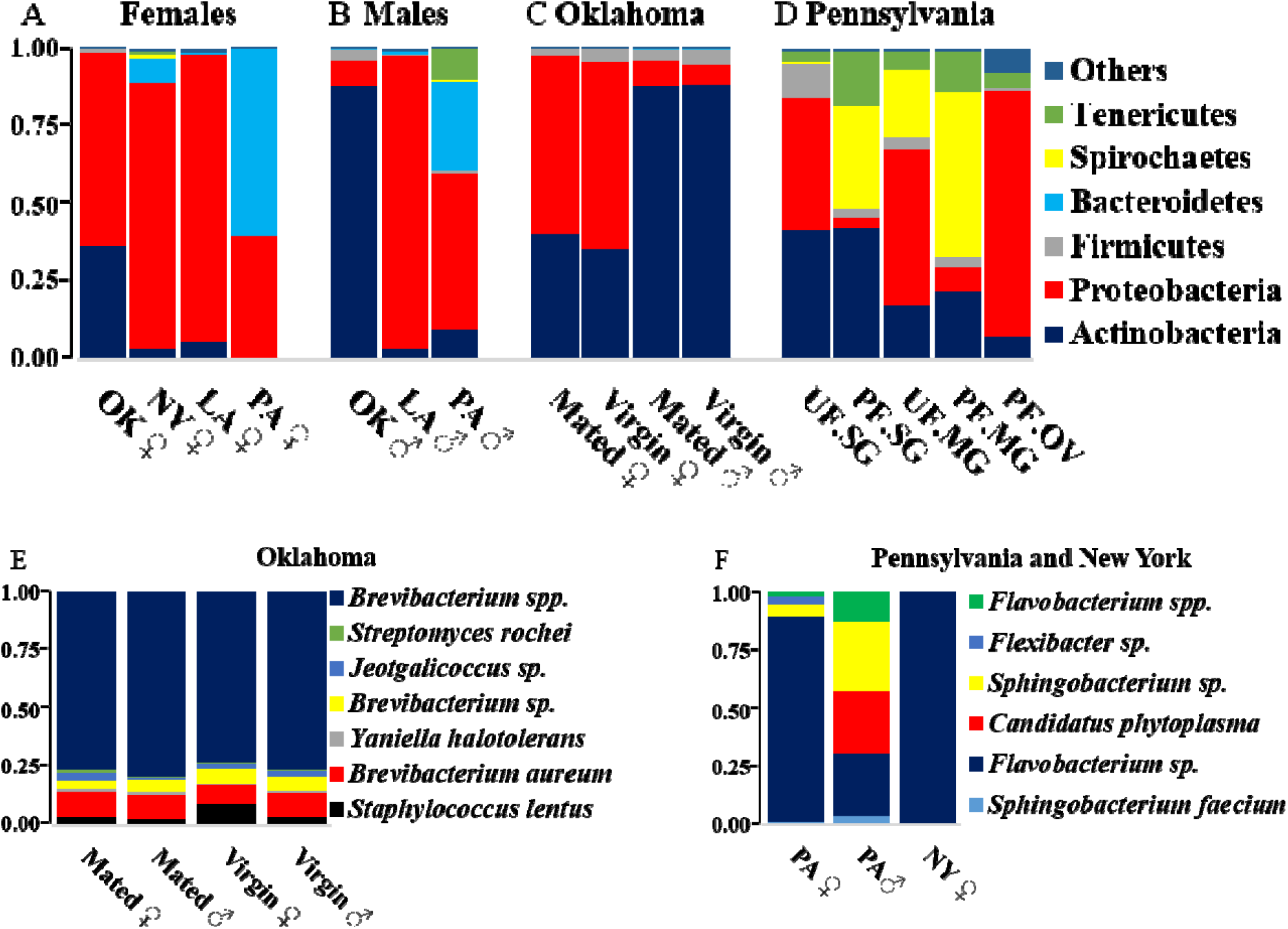
Bacterial profiling and relative abundance in *Ixodes scapularis* ticks at phylum level A) among female ticks B) Male ticks from Louisiana (LA), Pennsylvania (PA), Oklahoma (OK) C) and between virgin and mated ticks from Oklahoma D) in unfed and partially fed tick tissue procured from Pennsylvania (PA). E) at species level in Oklahoma ticks and **F) Pennsylvania ticks. ** If ticks from a particular location have shown significant difference in relative bacterial abundance at phylum level, have further been shown at species level

**Figure 1:**
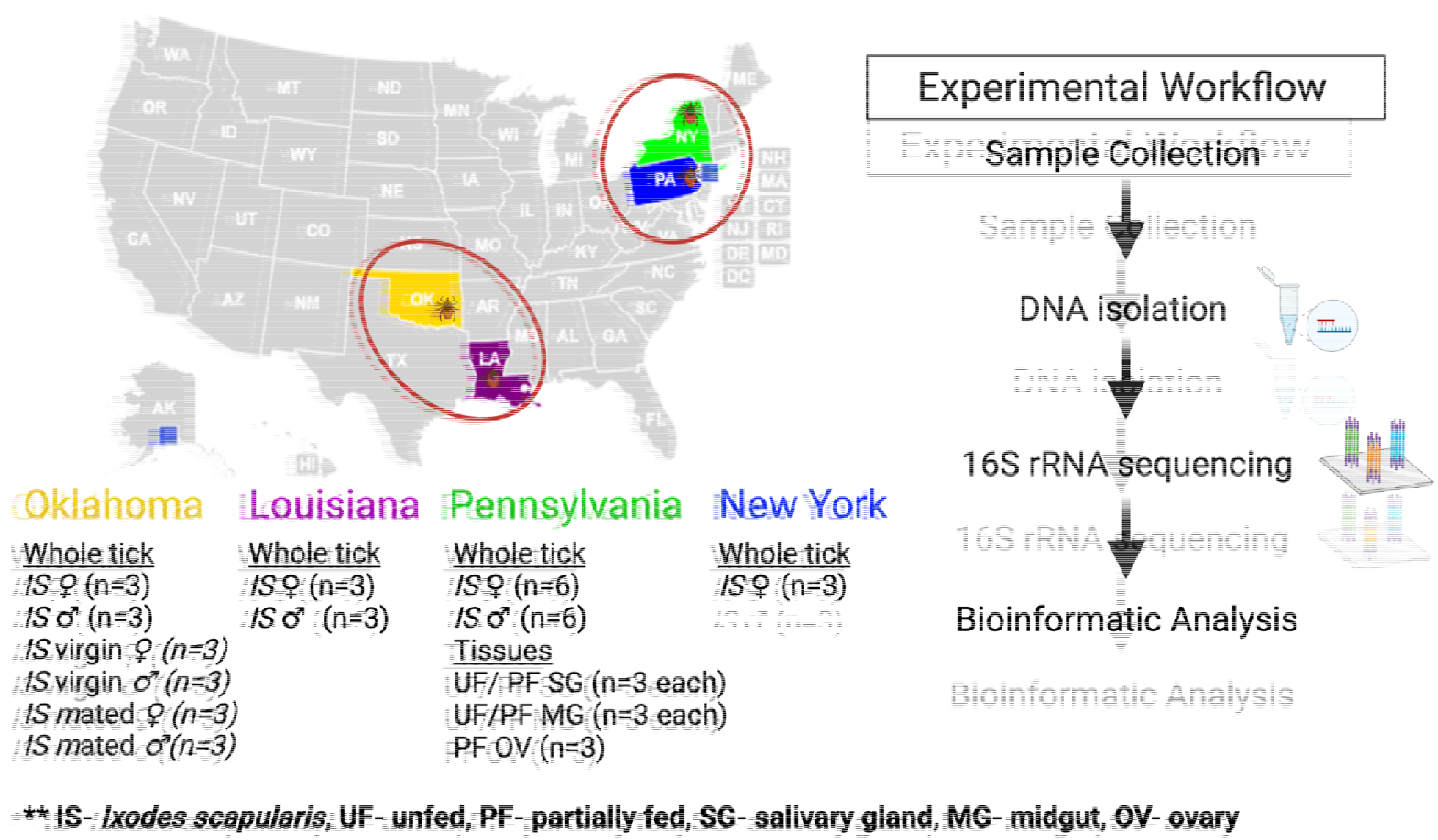
Schematic diagram of the microbiome study of the black-legged tick (*Ixodes scapularis*).

### 2.2 Comparative analysis of bacterial communities residing within Ix. scapularis

We observed substantial variation in bacterial profiles in tick populations from LA, NY, OK, and PA. Phyla Actinobacteria, Proteobacteria, Firmicutes, and Bacteroidetes cover for > 90 % reads for ♀ ticks from these populations (Supplementary Table S1). Proteobacteria were found to be the dominant phylum among all ♀ tick populations, excluding ♀ PA population, in which Bacteroidetes were dominant. A relative abundance of 36% Actinobacteria was present in ♀ OK ticks, but in other ♀ ticks, its coverage was substantially low as 0.58%, 6.13%, and 3.45% in PA, LA, and NY, respectively (Supplementary Table S1). Interestingly, ♀ PA ticks showed 60% relative abundance of Bacteroidetes, whereas this phylum was substantially less well represented in ♀ OK, NY, and LA, at 0.07%, 7.5% and, 0.49%, respectively (Table1). In ♂ ticks, a significantly high level of Actinobacteria (88%) was found in OK ticks, and levels were dramatically lower in LA and NY ♂, at 3.5%, 10%, respectively (Supplementary Table S1). NY ♂ ticks were not included in this study due to unavailability. The level of Proteobacteria was also found much lower in ♂ OK (6.19%) than ♂ LA (94.03%) and ♂ PA (50%) (Supplementary Table S1). Bacteroidetes (29%) and Tenericutes (10.4%) were substantially higher in ♂ PA than other whole unfed male ticks, ♂ OK (0.26% - Bacteroidetes), ♂ LA (1.49% - Bacteroidetes), while Tenericutes were rare in other ♂ ticks (Supplementary Table S1).

Bacterial profiling of mated and unmated tick populations were similar at phylum level (Figure 1C). Among these ticks, the dominant phylum in both ♂ and ♀ were Actinobacteria (∼88%) and Proteobacteria (∼56-60%) respectively (Figure 1A-B). Nonetheless, both ♂ and ♀ OK ticks contained ∼2-5% Firmicutes (Figure 3B-C), but other tick populations (NY & PA) contained either <1% or negligible Firmicutes (Figure 1A, 3B). Further analysis of OK ticks at species level showed Firmicutes’ presence, including *Jeotgalicoccus* sp. And *Staphylococcus lentus* (Figure 1E). At the same time, Actinobacteria included *Brevibacterium aureum, Yaniella halotolerans, Brevibacterium* sp., and *Streptomyces rochei* species (Figure 1E). As expected, both NY and PA tick populations were infected with a significantly higher percentage of Spirochetes, ranging from 2% reads in NY (♀) and 0.14% in PA (♂) respectively (Figure 1A-B). The prevalence of *B. burgdorferi* s. l. in *Ix. Scapularis* ticks from NY, and PA ranges from 40%- 70% [8-9]. All the ticks and tick tissues excluding ovary were infected with *B. burgdorferi* s.l. The results presented here are comparison of percentage *B. burgdorferi* s.l. reads among all microbial reads in the sample. For example, the tick from PA and NY are *B. burgdorferi* s.l. infected and 16S rRNA sequencing yielded a total of 100 reads from these ticks. Among those 100 reads, 2 and 0.14 reads belong to *B. burgdorferi* s.l. from NY and PA ticks, respectively. The rest of the reads belong to other microbes residing within these samples.

The PA ♂ and ♀ ticks contained ∼60% and 28.8% Bacteroidetes respectively and 7.48% in female ticks from NY (Figure 1A-B). Bacteroidetes included *Sphingobacterium faecium, Flavobacterium* sp., *Sphingobacterium* sp., *Flexibacter* sp. And *Flavobacter* spp. (Figure 3F). However, spirochetes were absent in both LA, and OK tick populations, and only ∼1.49% reads with Bacteroidetes were detected in the LA ♀ ticks (Figure 1 & Supplementary Table S1).

Differences and variations were also noted in bacterial composition of NY and PA ♀ tick populations. Proteobacteria (86%) was the dominant phylum in NY ♀ tick populations, whereas Bacteroidetes (60%) was dominant phylum in PA ♀ ticks (Figure 1A). Bacterial profiling of the NY ♀ and the OK ♀ ticks were significantly different at both phyla and species level (Figure 3A, Table.S7 species level profiling as supplementary data). The OK ♀ ticks contained 36.3%, 62.32%, 1.23%, and 0.06% reads of phylum Actinobacteria, Proteobacteria, Firmicutes, Bacteroidetes, and Spirochetes, respectively. However, the NY.♀ ticks harbor 3.45%, 85.75%, 0.28%, 7.48% and 1.96% reads of Actinobacteria, Proteobacteria, Firmicutes, Bacteroidetes and, Spirochaetes. Interestingly, both NY and OK tick populations at the species level demonstrated considerable differences in their bacterial profiling and abundance (Supplementary Table S2). Specifically, bacterial species such as *Brevibacterium* spp. (OK ♀ = 46.5%, NY ♀=0.33%), *Pseudomonas lurida* (OK ♀=0.17%, NY ♀ = 23.4%), *Brevindumonas diminuta* (OK ♀=0.05%, NY ♀= 8.42%), *Rickettsia* spp. (OK ♀=29.23%, NY ♀= 15.8%), *Flavobacterium* spp. (OK ♀=0.04%, NY ♀=6.62%), *Stenotrophomonas rhizophila* (OK ♀=0.0358, NY ♀=14.52%), *Pseudomonas fluorescens* (OK ♀=0.04%, NY ♀=8.16%), *Brevibacterium aureum* (OK ♀=6.22%. NY ♀=0.04%) demonstrated substantial differences in bacterial profiling between these female ticks collected from different geographical regions, specifically the northeastern and southern United States.

### 2.3 Bacterial composition of unfed and partially engorged tick tissues from Pennsylvania (PA)

An important difference in phylum-level microbial profiling was detected among different tissues collected from unfed and partially blood-fed ♀ tick tissues (Figure 3D, Supplementary Table S3). There was substantial difference in relative abundance of Proteobacteria (UF.SG – 42.27%, PF.SG-3.43%), Firmicutes (UF.SG – 11.18%, PF.SG-3.29%) and Spirochetes (UF.SG – 0.26%, PF.SG-32.78%) between unfed (UF) and partially fed (PF) salivary gland (SG) tissues (Figure 3D, Supplementary Table S3). In midgut (MG) tissues, trend for Proteobacteria (UF.MG – 50.34%, PF.MG-7.7%) and Spirochete abundance (UF.MG – 21.6%, PF.MG-52.7%) were similar to that of salivary gland (SG) (Figure 1D, Supplementary Table S3). Unfed ovary was not available for this work. In the case of the partially fed ovary (PF.OV), the level of Proteobacteria (85.6%) was considerably higher than other tissues while Spirochete level (PF.OV – 0.12%) was substantially low (Figure 1D, Supplementary Table S3). It was noted that Bacteroidetes were the most dominant (Relative abundance, 60%) phylum of PA female ticks at unfed whole tick level but were completely absent at tissue level (SG, MG, OV).

A considerable difference in microbial profiling at phylum level was detected among different tissues collected from the PA unfed and partially blood-fed ♀ tick tissues (Figure 1D). Unfed salivary glands contained 42%, 42.27%, ∼11.2% respectively of Actinobacteria, Proteobacteria, and Firmicutes, whereas partially blood-fed salivary glands harbored 42% Actinobacteria reads followed by a substantially reduced level of Proteobacteria (3.43%) and Firmicutes (3.2%). Interestingly, both Spirochetes and Tenericutes levels increased from 0.26% to 32.77% and 3.37% to 17.66%, respectively (Figure 1D, Supplementary Table S3). Table 3 shows the bacterial species abundance in tick tissues. Unfed midgut contained 50.34%, 21.61% and 17.5% Proteobacteria, Spirochetes, and Actinobacteria, respectively, while partially engorged midgut of Proteobacteria decreases substantially to 7.7%, Spirochetes increases drastically to 52.7%. The unfed ovary was unavailable, and the partially engorged ovary contained 85.56% of Proteobacteria, 8.11% of Actinobacteria, 5.24% of Tenericutes, and <1% of Firmicutes and Spirochetes (Supplementary Table S3).

### 2.4 Presence of Rickettsia buchneri in partially fed tick tissues and pattern of likely competitive interactions with B. burgdorferi s.l

*R. buchneri* reads in partially blood-fed salivary glands decreased drastically to 0.33% compared to unfed salivary glands (35.38%); however, levels of *B. burgdorferi* **s.l**. increased from 0.23% to 32.77% (Figure 2A). Similarly, partially engorged midgut tissues also showed the same trend as salivary glands, i.e., the level of *Borrelia* increased substantially, and rickettsia level decreased substantially (Figure 2B, Supplementary Table S4). Interestingly, the partially-fed ovaries showed ∼85% reads of rickettsia and, as expected, 0.12% of *Borrelia* since it is not vertically transmitted to the next generation (Figure 2B and Supplementary Table S4). These results demonstrated the infection of *R. buchneri* in the midgut (UF / PF: 46% / 4%) and salivary glands (UF/PF: 35%/0.3%), and these levels highlight the substantial reduction in the level of *R. buchneri* upon blood feeding. Mean absolute abundance of *B. burgdorferi* s. l. (UF.SG = 21.67, PF.SG = 4820.67, UF.MG = 3875.33, PF.MG = 16284.70, PF.OV = 54) and *R. buchneri* (UF.SG = 7154.3, PF.SG = 44, UF.MG = 11044.67, PF.MG = 292.67, PF.OV = 38828) also indicate the same trend as that of their relative abundances (Figure 2C, Supplementary Table S5).

**Figure 2.**
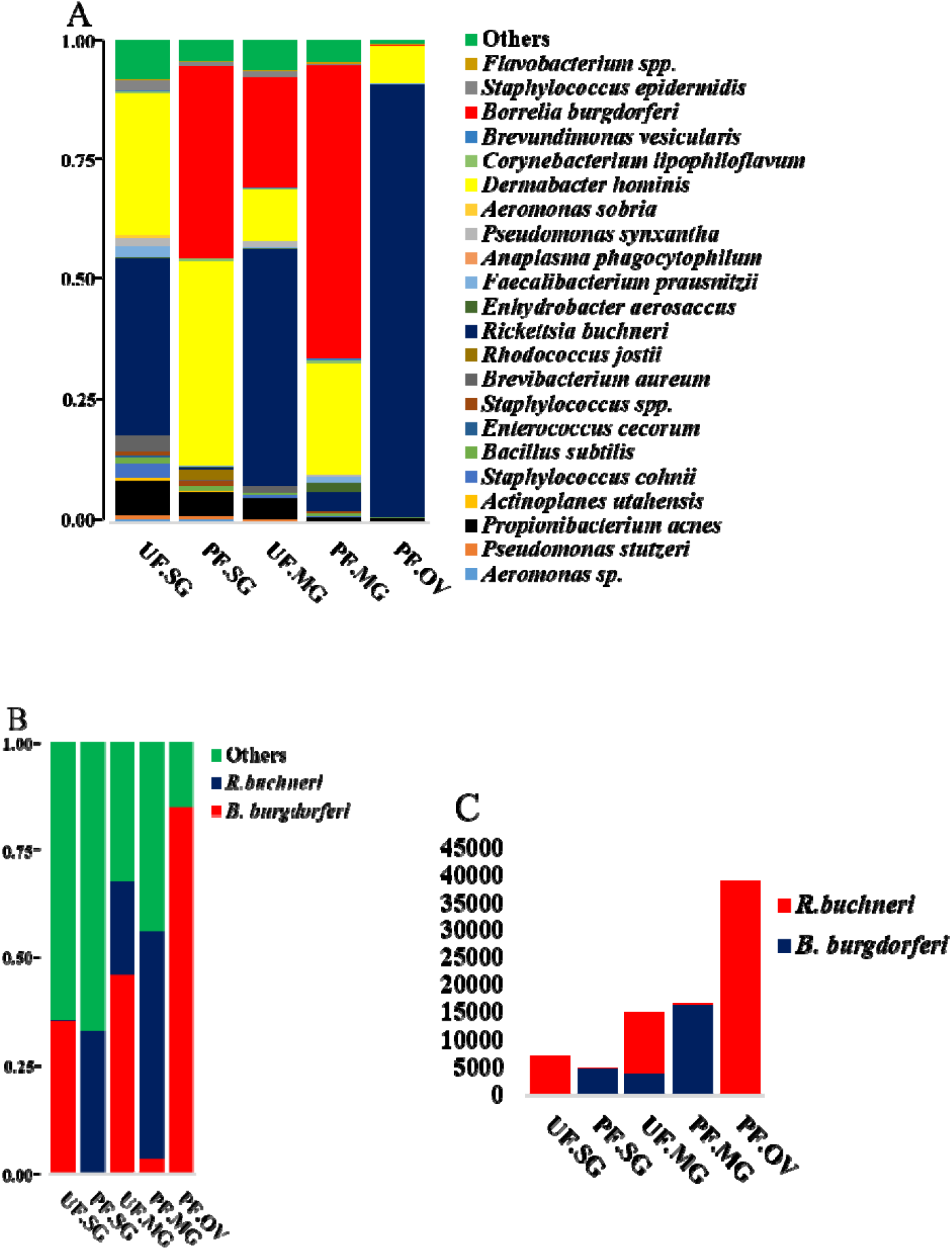
Microbial profiling at A) species level in unfed and partially fed tick tissues (SG, MG, OV). These field-collected *Ixodes scapularis* ticks were procured from Pennsylvania. B) Probable competitive interaction between endosymbiont *Rickettsia buchneri* and *Borrelia burgdorferi*. UF-unfed, PF-partially fed, SG-salivary gland, MG-midgut, OV-ovary. C) Absolute abundance (mean of the number of sequences of biological replicates) of *Borrelia burgdorferi* and *Rickettsia buchneri* in tick tissues from Pennsylvania (PA). UF-unfed, PF- partially fed, SG-salivary gland, MG-midgut, OV-ovary.

### 2.5 Diversity analysis

Kruskal-Wallis non-parametric tests were performed to determine the effects of tick geographic location (LA, PA, NY, OK) on α-diversity metrics using QIIME 2. Faith_pd- diversity index, p-value and adjusted-p-value (q-value) between the samples are as follows: LA and PA ♀ ticks (H=5.4, p-value = 0.02, q-value=0.101), LA ♂ ticks and OK mated ♂ ticks (H=3.85, p-value=0.049, q-value=0.101), NY and PA ♀ ♂ ticks (H=5.4, p-value=0.02, q-value=0.101), OK and PA ♀ ticks (H=5.4, p-value=0.02, q-value=0.101), and OK and PA virgin ♂ (H=4.5, p-value=0.0338, q-value=0.101) (Figure 3C). PA ticks were phylogenetically least rich while OK virgin ♂ are the richest in bacterial diversity among samples studied here. At the tissue level, midgut from partially engorged ticks were found the richest in diversity and ovaries were the least rich in bacterial diversity (Figure 3A). Tissue samples did not demonstrate statistically significant evenness (pielou_e) in bacterial diversity (Figure 3B). Mated and virgin OK ♂ ticks contain similar bacterial profiles (Figure 3D).

**Figure 3.**
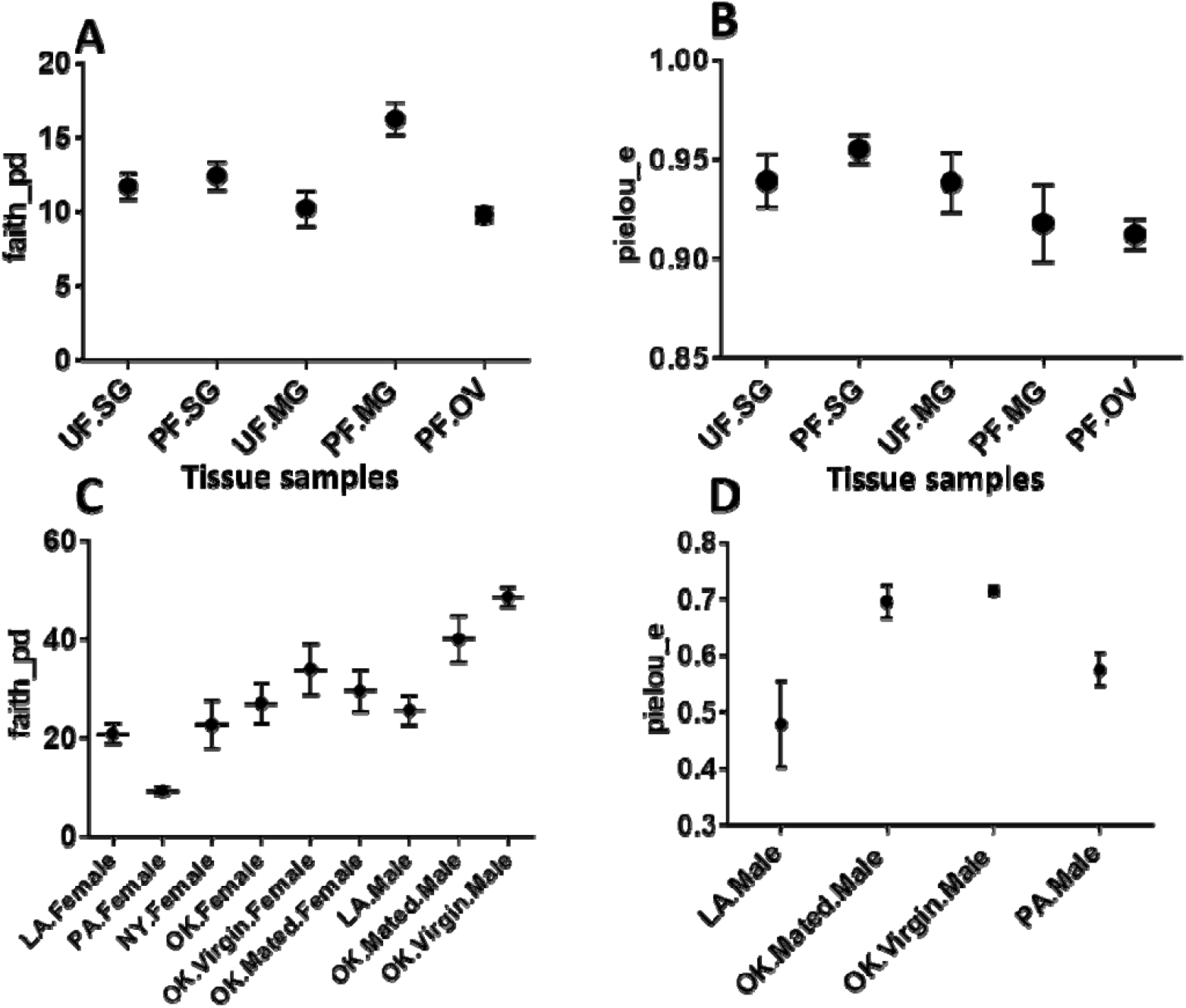
Comparisons of bacterial alpha diversity among tick tissue samples (A, B) and ticks collected from different geographical locations (C, D). Plotted data represent alpha-diversity based on Faith’s phylogenetic distance (faith_pd) demonstrating relative richness among bacterial communities, while Pielou’s index represent the evenness of the bacterial community among different samples. Upon Benjamin & Hochberg correction, all the adjusted-p-values (q-values) were insignificant. UF-unfed, PF-partially fed, SG-salivary gland, MG-midgut, OV-ovary, LA-Louisiana, PA-Pennsylvania, NY-New York, OK-Oklahoma.

As it is clear from the Principal Coordinate analysis (PCoA) plot of unweighted unifrac distances, most of the tick samples from different locations (NY, OK, PA and LA) contained different bacterial profiles (p=0.001) as determined by PERMANOVA (Permutational multiple analysis of variance). PCoA of unweighted UniFrac distances of bacterial communities showed that the first two axes (Axis1 and Axis2) explained 31.14% and 8.5% of the variation in the data respectively (Figure 4B). PERMANOVA analysis of Unweighted UniFrac distances, p-value and adjusted p-values (q-value) in several of the samples when compared pairwise were as follows LA and PA ♀ ticks (pseudo-F= 7.33, p-value= 0.011, q-value = 0.095625), LA.♂ and PA.♀ ticks (pseudo-F= 6.10, p-value=0.017, q-value = 0.095625), NY.♀ and PA.♀ (pseudo- F=7.19, p-value= 0.013, q-value = 0.095625), OK.♀ and PA.♀ (pseudo-F= 10.80, p-value= 0.008, q-value = 0.095625), OK Mated.♀ and PA♂(pseudo-F= 2.99, p-value= 0.047, q-value = 0.147), OK.Mated.♂ and PA.♂ (pseudo-F= 3.20, p-value= 0.03, q-value = 0.135), OK.Virgin♀and PA.♀ (pseudo-F= 8.124, p-value= 0.017, q-value = 0.095625), OK.Virgin.♂ and PA♂(pseudo-F= 3.76, p-value= 0.027, q-value = 0.135) (Supplementary Table S6). For the unweighted Unifrac of whole tick samples, OK tick samples (Mated ♀, Mated ♂, Virgin.♀, Virgin.♂), LA ticks (♂, ♀), and PA (♀), and PA (♂) are making separate clusters for the most part. Interestingly, it also shows few outliers, such as PA ♂ and NY ♀ (Figure 4A). Unweighted Unifrac PCoA plot (Figure 5B) showed that partially fed midgut is distinct from unfed midgut at the tissue level. In this case, PCoA of unweighted UniFrac distances of bacterial communities in the first two axes (Axis1 and Axis2) explained 30.5% and 13.27% of the variation in the data. Again in the weighted Unifrac PCoA plot (Figure 5A) also, partially fed midgut (PF.MG) has clustered distinctly than unfed midgut (UF.MG) and other tissues indicating that the dominant bacterial diversity in the partially fed midgut is distinct than other tissues. All other tissues except partially-fed midgut have clustered together separately, indicating common dominant bacteria.

**Figure 4.**
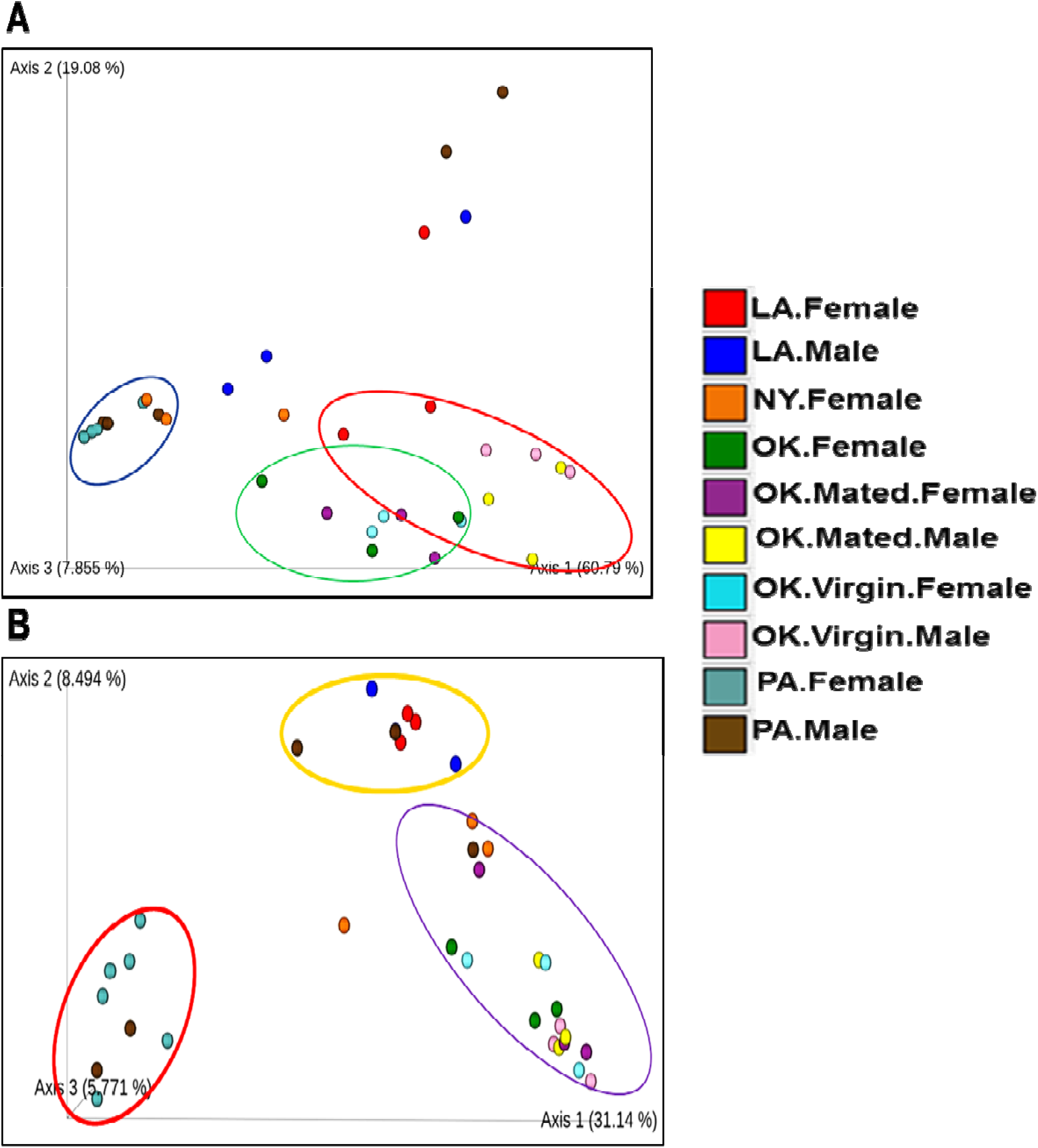
PCoA plot for male and female ticks collected from OK, LA, NY and PA. A) Weighted UniFrac distance, the bacterial composition in female from Pennsylvania (PA. Female), New York (NY. Female) and male from Pennsylvania cluster together. B) Unweighted unifrac plot.

**Figure 5.**
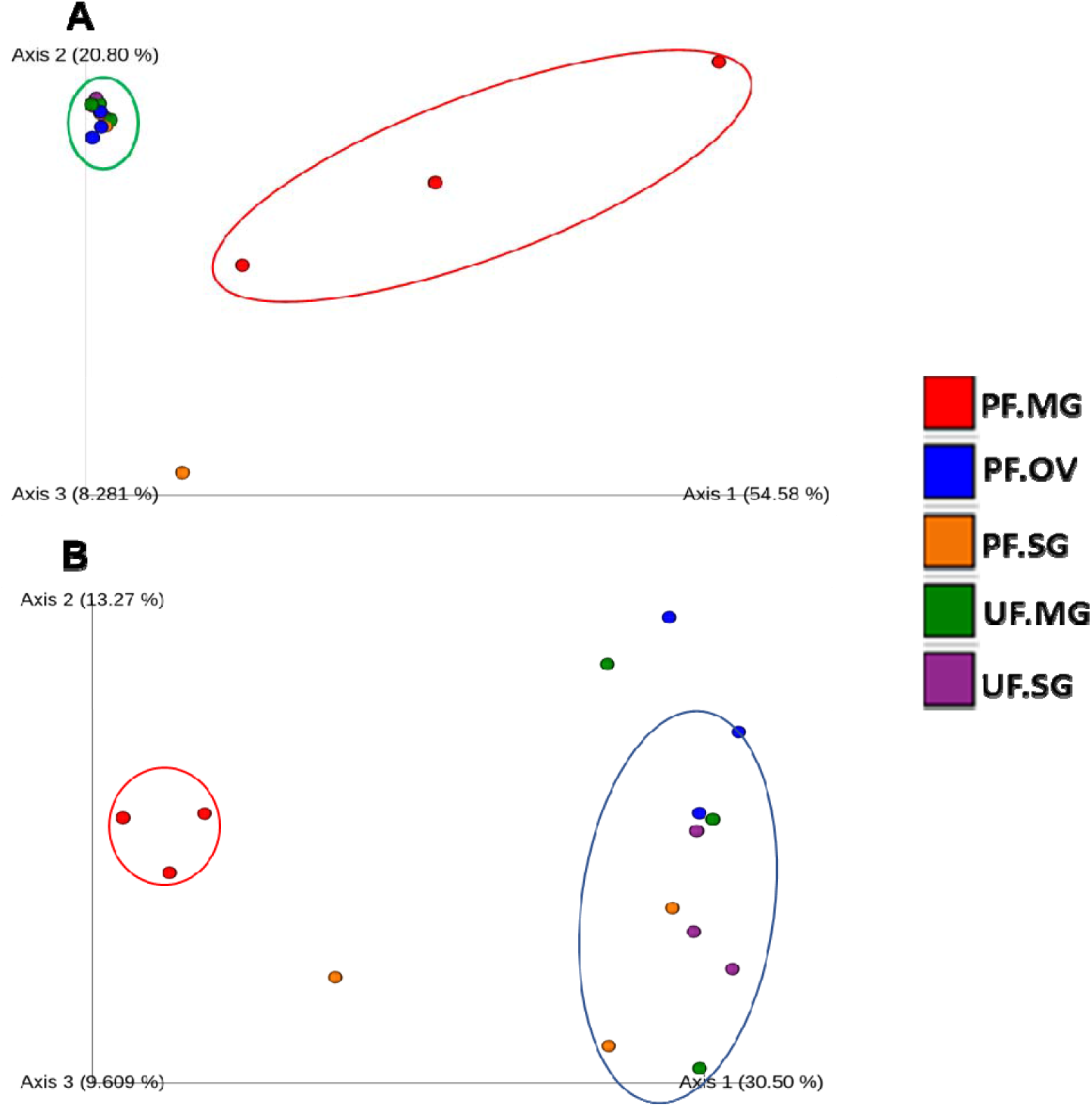
PCOA plot for unfed and partially fed tick-tissues collected from Pennsylvania. A) Weighted Unifrac B) Unweighted Unifrac. UF-unfed, PF-partially fed, SG-salivary gland, MG-midgut, OV-ovary.

### 2.6 Microbial Interactions

Network analysis of microbial interactions using the SparCC permutations revealed 726 interactions amongst 36 taxa when applied to life stage dataset from all regions, out of which 358 were positive interactions while 360 were negative interactions (Figure 6; Supplementary table S7). All identified taxa belonged to the phylum Nanoarchaeaeota (Unidentified genus), Proteobacteria (18), Spirochates (1), Actinobacteria (8), Bacteroidetes (6), and Firmicutes (2). Most interactions were seen with the bacteria in the genus *Flavobacterium, Pseudomonas*, and *Brevibacterium*. Bacteria genus from pathogenic group belonging to *Rickettsia* and *Borreliella* were also identified in the network interactions with *Rickettsia* interacting with more members compared to *Borreliella*.

**Figure 6.**
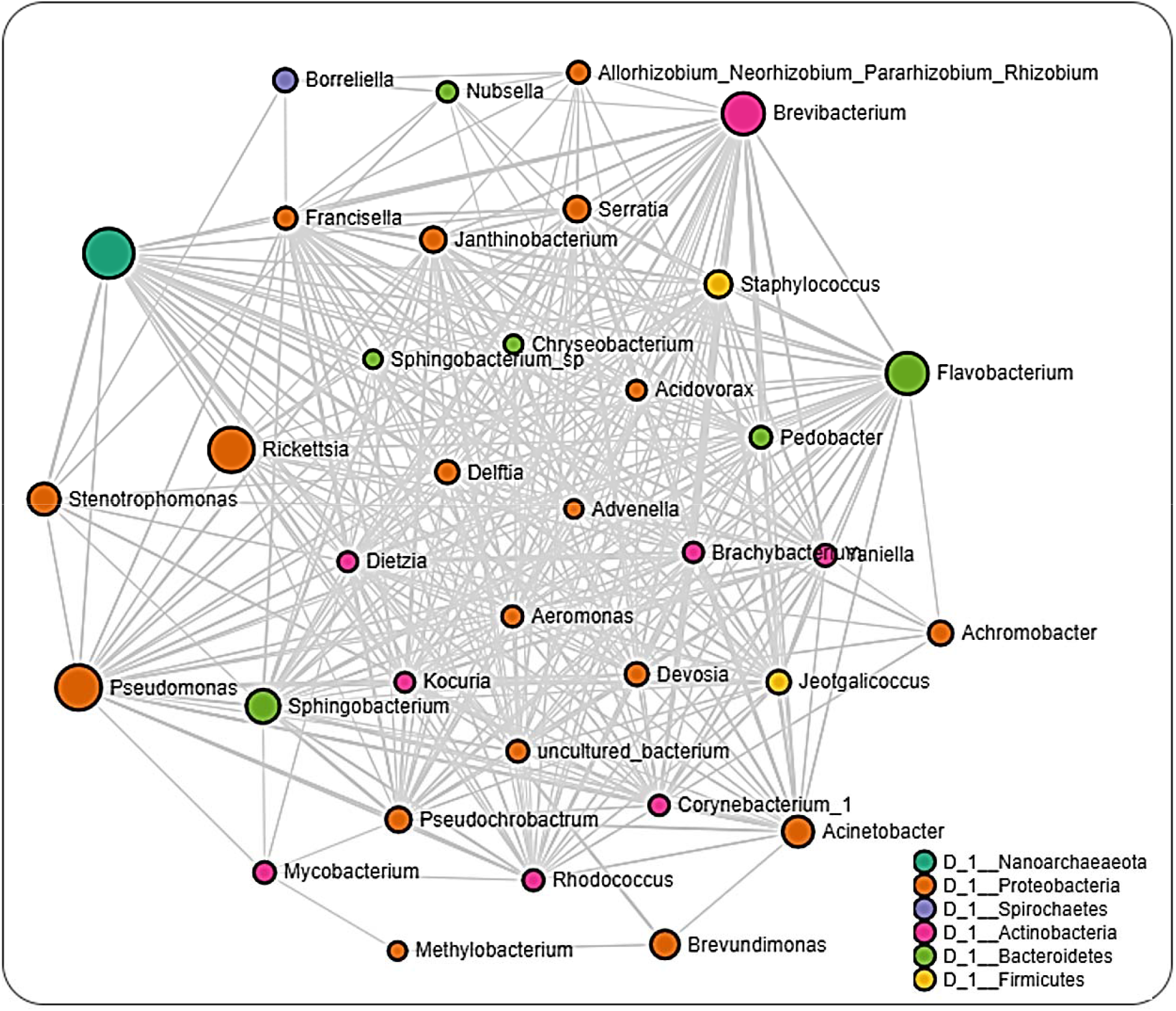
*Correlation network analysis*. Correlation network maps generated using the SparCC approach. Correlation network with nodes representing taxa at the genus level and edges representing correlations between taxa pairs. Node size correlates to the number of interactions a taxon is involved with. Color coded legend shows the bacteria phylum each taxon belongs to.

The interaction cluster in which *Rickettsia* was detected revealed several positive and negative correlations notably positive correlation with *Francisella*. In the network, *Borreliella* was identified in the same cluster as *Allorhizobium_Neorhizobium_Pararhizobium_Rhizobium, Nubsella, Stenotrophomonas* and notably *Francisella* all of which were positively correlated to one another. The network analysis performed on dataset from the different regions showed that ticks from Pennsylvania had the most significant partial correlations of 140 involving the most OTUs (32) (Figure 7; Supplementary table S8). *Borreliella* was identified across the datasets but was observed to be correlated with different bacteria genera from each region. *Borreliella* from Pennsylvania dataset was negatively correlated with *Brevundimonas* and positively correlated with *Jeotgalicoccus* (Supplementary table S8).

**Figure 7.**
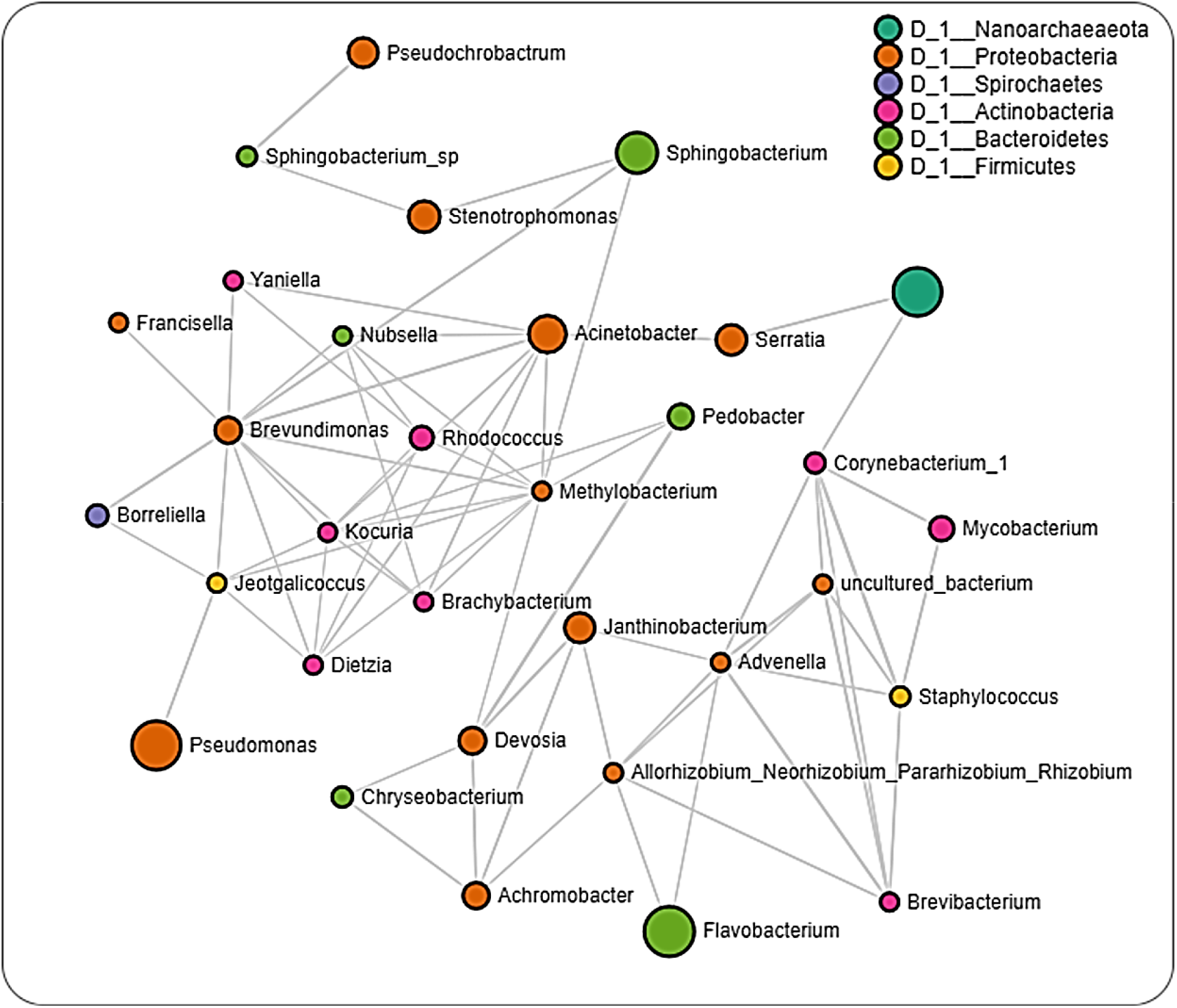
*Correlation network analysis across ticks from Pennsylvania*. Correlation network maps generated using the SparCC approach. Correlation network with nodes representing taxa at the genus level and edges representing correlations between taxa pairs. Node size correlates to the number of interactions a taxon is involved with. Color coded legend shows the bacteria phylum each taxon belongs to.

In ticks from Louisiana, we observed negative partial correlations between *Borreliella* and *Delftia*, and positive partial correlations between *Borreliella* and *Rickettsia*, while dataset from Oklahoma only showed partial negative correlation between *Borreliella* and *Brevundimonas* (Figure 8 and 9; Supplementary table S9 and S10). We however did not detect the presence of *Rickettsia* in the dataset from Pennsylvania ticks. Datasets from Louisiana and Oklahoma both had *Rickettsia* in their network interactions and interestingly shares similarity in their partial positive correlations with the genus *Advenella* and *Sphingobacterium*. While datasets from both Oklahoma and Louisiana contained *Borreliella* and *Rickettsia*, a direct significant correlation was only observed between both bacteria in the dataset from Louisiana.

**Figure 8.**
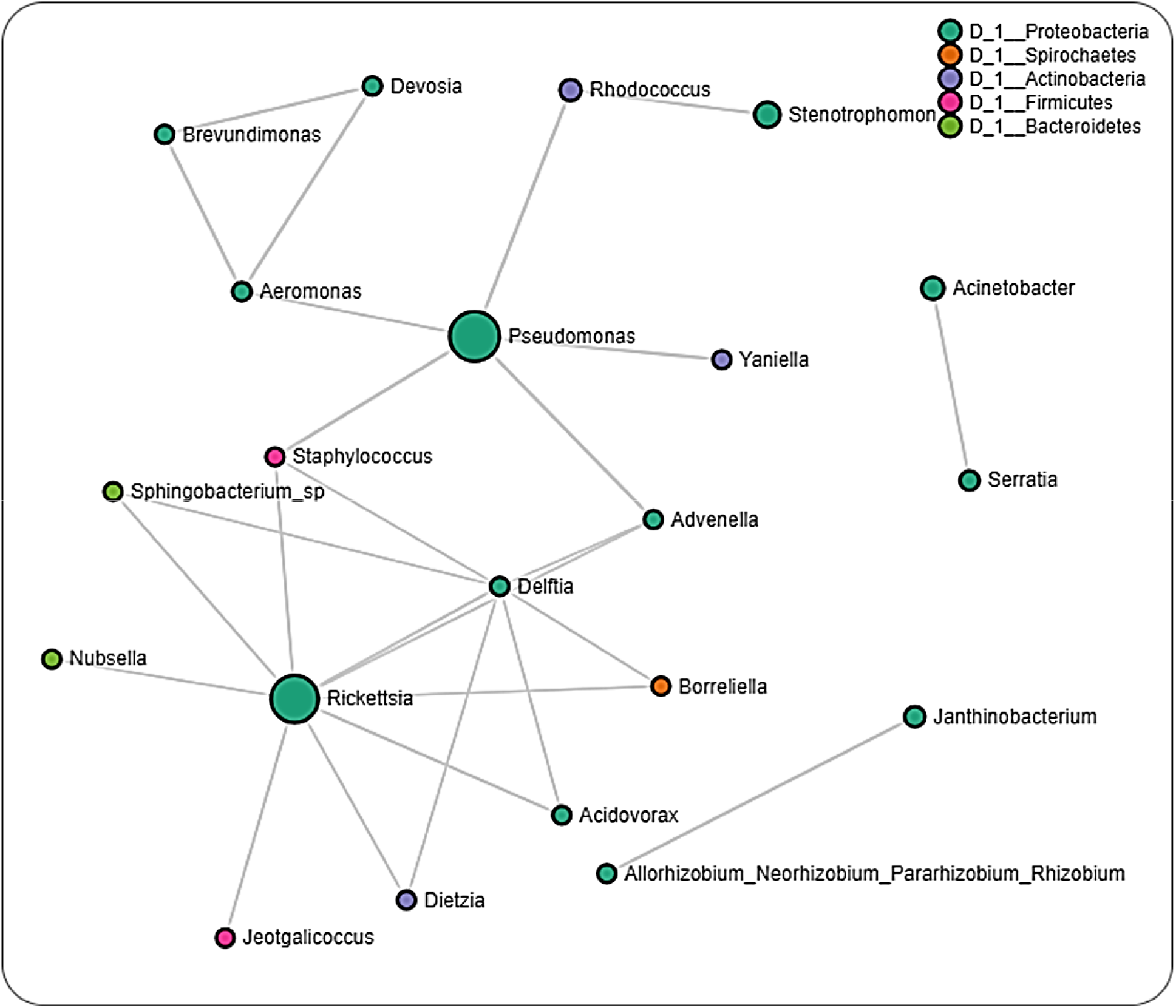
*Correlation network analysis across ticks from Louisiana*. Correlation network maps generated using the SparCC approach. Correlation network with nodes representing taxa at the genus level and edges representing correlations between taxa pairs. Node size correlates to the number of interactions a taxon is involved with. Color coded legend shows the bacteria phylum each taxon belongs to.

**Figure 9.**
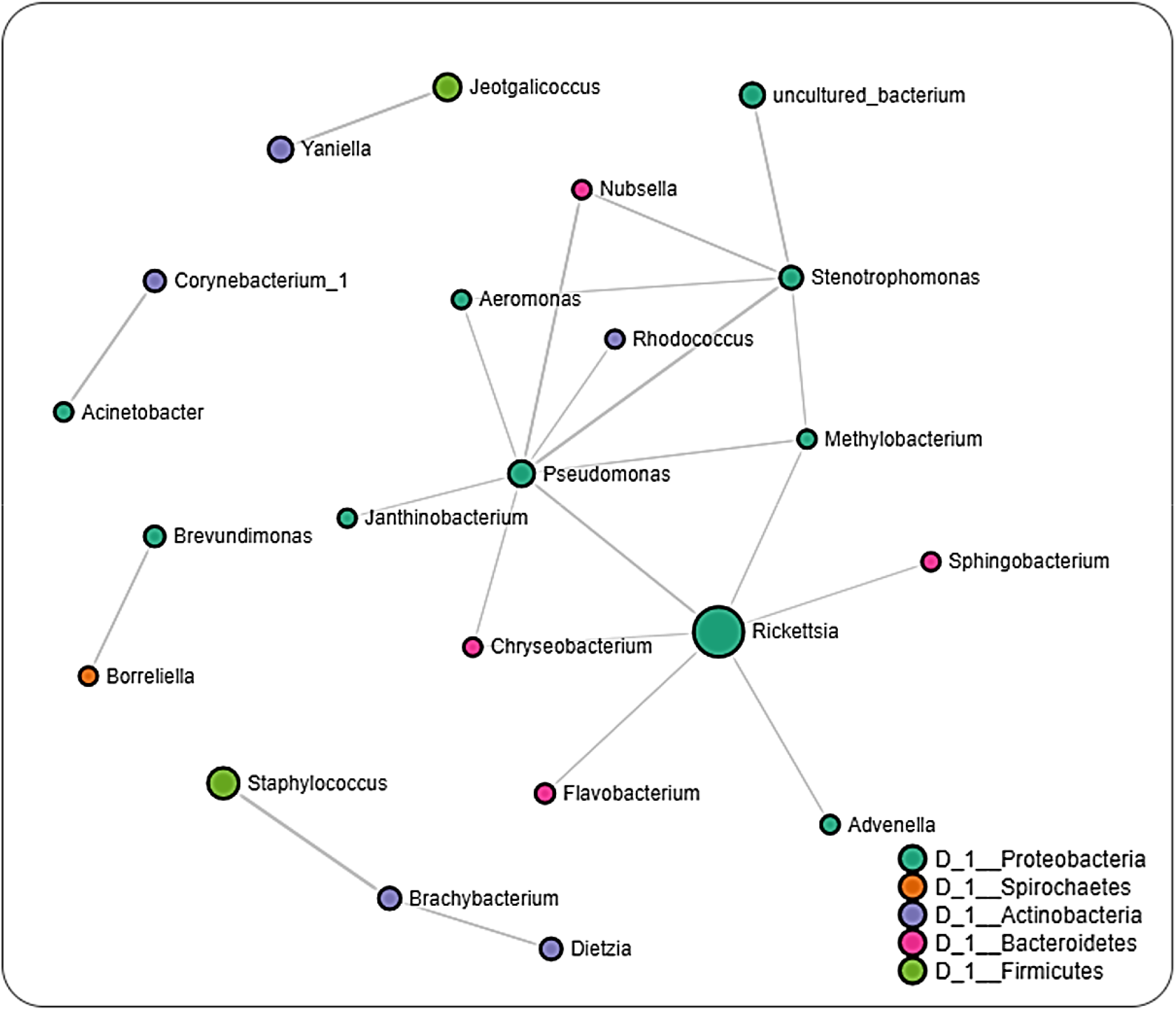
*Correlation network analysis across ticks from Oklahoma*. Correlation network maps generated using the SparCC approach. Correlation network with nodes representing taxa at th genus level and edges representing correlations between taxa pairs. Node size correlates to th number of interactions a taxon is involved with. Color coded legend shows the bacteria phylum each taxon belongs to.

## 3. Discussion

In this study, the microbiome of *Ix. scapularis* from Lyme disease-endemic (NY & PA) and non-endemic (LA & OK) regions showed substantial variation by both geography and sex at the organismal and tissue levels. A variation in the microbiota of *Ixodes* ticks with regard to geography and sex has been suggested [14]. Our results are not in complete coherence with other studies; [31- 34] as those studies were conducted under a variety of different conditions to examine tick’s microbiome. For example, Sperling et al., [34] analyzed the ticks collected off the cats and dogs from Alberta (Canada), whereas Kwan et al., [31] conducted their studies on *Ixodes pacificus* ticks. Several studies have also identified variation in microbiome among different tick species [35], sex [36-37], and geography [16, 14]. It remains unclear what factors drive the microbiome’s natural variation [33]. The impact of host and environmental drivers on the microbiome composition is still an under-investigated area [33]. Our study showed the dominance of Proteobacteria in most of the tick samples except PA ♀ and OK ♂ tick samples. Firmicutes were found only in OK ticks, and Bacteroidetes only in NY and PA ticks. These results raised an important question of whether these ticks in different geographic regions maintain a distinct microbiome and whether these variations among the microbial composition and communities are driven by the host animals or even by soil bacteria [38].

Our data also showed comparatively higher *B. burgdorferi* s.l. reads (∼2.5%) in the NY and PA female ticks compared to LA and OK tick populations [6, 4]. *B. burgdorferi* s.l. reads were also detected in the PA male ticks (Figure 1B), which is surprisingly higher than other male tick samples. Bacteroidetes were dominant in the PA female tick, whereas in both unfed and partially engorged tick tissues, the level was surprisingly low (Figure 1A, B, and D). It suggests bloodmeal-induced changes in the microbiome composition in tick tissues. Interestingly, studies pointed out that vertebrate hosts do not influence the bacterial composition within adult flea and tick species (*Dermacentor variabilis* and *Ix. scapularis*) [35], but blood-feeding in immature developmental stages significantly impacts the bacterial community structuring [39]. Before and immediately after blood feeding of *Ixodes persulcatus* on rats, a report showed similar alpha-diversity but significantly different bacterial profiling of tick [37].

In the present study, the endosymbiont *R. buchneri* was found in the salivary gland, midgut, and ovary, and its level drops significantly when *B. burgdorferi* s.l. multiplies in tick tissues upon blood-feeding (Figure 2B), suggesting a possible competitive interaction that is purely speculative at this point and needs more work to support it. Although absolute abundance data of *R. buchneri* and *B. burgdorferi* s.l. (Figure 3 and Supplementary Table S5 supports this possible competitive interplay. Recent elegant work from Oliver et al., [40] has shown the growth dynamics of *R. buchneri* in *Ix. scapularis* ticks. This type of competitive interaction has also been demonstrated between *B. burgdorferi* s.l. and gut microbiota in the midgut of *Ix. Scapularis* [41] and by *Rickettsia* spp. in the ovary of *Dermacentor andersoni* [42]. The mechanism of co- existence of *B. burgdorferi* s.l. with other microbial pathogens inside ticks remains unknown [43] and is an area of active research. As *B. burgdorferi* s.l. is extracellular and *R. buchneri* is an intracellular microbe, they might not have physical interaction but they may affect colonization by activating immune and reactive oxygen species pathways [43-45]. The endosymbiont *R. buchneri* is also known as “rickettsial endosymbiont of *Ix. scapularis*” [46]. The genome of *R. buchneri* is significantly larger (> 2Mb) than that of pathogenic rickettsiae, and it also has the capability to synthesize vitamin B components, biotin, and *de novo* folate [46]. As indicated by previous studies, one of the major roles of the symbionts from ticks and other obligate hematophagous arthropods is to provide vitamin B, that ticks are deficient due to their exclusively sanguineous diet [47-48]. The presence of *R. buchneri* was reported to be restricted to tick ovaries [49-50], but a recent study has reported its colonization into tick salivary glands [48]. It might evolve as a pathogen as it fulfills the prerequisite for an endosymbiont to be transmitted to the vertebrate host by getting colonized into tick’s salivary glands [48]. Our data has shown the colonization of this rickettsial species into salivary glands (UF.SG-35.38%, PF.SG-0.33%) and midgut (UF.MG-46.09%, PF.MG-3.52%) during unfed stage, but the level of *R. buchneri* reduces significantly after blood feeding. It is not necessary that this process would lead to a condition of pathogenicity, it might simply be a transmission route to other ticks through co-feeding. Nevertheless, a symbiont in the salivary gland might also be exposed to the host immune system, leading to an antibody response or can interact with pathogens in the salivary gland and may facilitate or impede their transmission [48]. Studies have shown that several genera of bacteria such as *Rickettsia, Coxiella, Francisella, and Midichloria*, persist transtadially and later get transmitted transovarially as a regular process [47, 51}. These endosymbionts might remain restricted to the arthropod host, sometimes may be transmitted to the vertebrate host or sometimes may cause disease [51, 47]. The distribution of reads for *R. buchneri* in the various tissues is consistent with the rest of the literature [48].

Network analysis across all life stage datasets from all regions revealed relatively equal number of positive (∼49%) and negative (∼51%) interactions to be present. Positive interactions between different bacteria taxa could indicate shared functionality, or even a shared niche within the host organism [52-54], whereas negative interactions would point towards an existing or potential competition between bacteria taxa. This would suggest that the microbiome of *Ix. scapularis* favors a balanced distribution between bacteria with potential synergistic and antagonistic interactions. This observation contrasts the detection of greater than 97% positive interactions in the *Ix. ricinus* microbiota as recently reported by Lejal et al. [55], thus indicating differences exist in the microbial-microbial interaction even between the same tick genera. Most of the interactions observed in the whole dataset were driven by bacteria belonging to non-pathogenic genera as indicated by the presence of *Flavobacterium, Pseudomonas* and *Brevibacterium* further suggesting a contribution of non-symbiotic commensal microbes to the overall microbiome of the *Ix. scapularis* ticks.

Interestingly, we identified OTUs belonging to the pathogenic *Rickettsia* and *Borreliella* genus from the network analysis, with *Rickettsia* observed to interact with more bacteria genera compared to *Borreliella*. Positive correlation was seen to exist between *Francisella* and *Borreliella*. The interaction between *Rickettsia* and *Borreliella* was also observed to vary across different geographical location. While our network analysis was carried out down to the genus level, the bacterial profile identified the *Rickettsia* identified to the species level as *R. buchneri*, the major endosymbiont of *Ix. scapularis*. The genus *Francisella*, which are endosymbionts, was significantly correlated with *Rickettsia* suggesting an indication of a co-dependency on two endosymbionts by *Ix. scapularis*. This hypothesis is in accordance with a recent report of a dual endosymbiont dependency observed between *Midichloria* and *Francisella* symbionts in *Hyalomma marginatum* ticks driven by a nutritional adaptation [56]. It has also been shown that, although, *R. buchneri* do possess the essential vitamin synthesis genes, some *Ix. Scapularis* harbors this endosymbiont that lacks these vitamin synthesis pathways indicating a non-obligatory or facultative endosymbiotic relationship [57-59]. This could explain the need to harbor a different class of endosymbiont in *Francisella* that would relieve the exclusive dependence on *R. buchneri* as seen in this study.

The detection of multiple endosymbiont species has also been identified in other tick species such as in the *Amblyomma maculatum* tick although the functional contribution to the tick biology has not been described [60]. *Rickettsia* and *Borreliella* were not identified within the same cluster in the network analysis of the whole dataset; however, both bacteria genera interacted differently in ticks from different locations. A much surprising observation was the presence of *Borreliella* and corresponding absence of *Rickettsia* from the network analysis on datasets of ticks from Pennsylvania. While the small size of the tissue dataset from these regions prevented us from carrying out network analysis, this observation was further supported by the exclusive presence of *R. buchneri* in unfed salivary glands and partially fed ovarian tissues. A shared presence of both bacteria in partially fed salivary glands and midgut tissue suggests a tissue driven microbial interaction. This observation contrast with the report of Aivelo et al. who reported positive correlations between Lyme borreliois *Borrelia* group, *Borrelia miyamotoi* and *Rickettsiella* in the sister tick *Ix. ricinus* suggesting a specific interaction dependent on the host tick specie.

## 4. Materials and Methods

### 4.1 Ticks

Only adult ticks were field collected from Louisiana (LA), New York (NY), and Oklahoma (OK) and, Pennsylvania (PA). Unfed adult *Ix. scapularis* ticks mate before attachment on a vertebrate host. Ticks were identified using standard morphological keys. To test the microbes’ transfer during mating of male and female unfed ticks (♂ & ♀) were used for these experiments. Unmated ♀ & ♂ ticks were kept in a vial for 48 hours and allowed to mate with their partners. Unfed adult ticks from PA were blood-fed as described earlier [61], and tissues were dissected from partially engorged female ticks. The dissecting solution was ice cold 100 mM 3-(N- Morpholino-propanesulfonic acid (MOPS) buffer containing 20 mM ethylene glycol bis-(β-aminoethyl ether)-N, N, N’, N’ -tetraacetic acid (EGTA), pH 6.8. After removal, salivary glands, midgut, and ovaries were washed gently in the same ice-cold buffer. The dissected tissues were stored immediately after dissection in DNA lysis buffer before isolating DNA. The experimental plan for this study is illustrated in Figure 1. Briefly, at least three biological replicates were used for each biological sample (unfed and partially engorged ticks) and tissue types (Midgut, Salivary glands, and ovary) from PA ticks. Additionally, three biological replicates for each of dissected tissues and whole tick were used in this study. The unfed ovary tissues were excluded from PA in these analyses due to the small size and technical challenge in getting enough DNA from a single dissected ovary. Partially engorged ovaries from PA ticks were dissected for these experiments.

### 4.2 DNA isolation

Genomic DNA was extracted from 1) individual whole unfed ticks from LA, NY, OK and, PA, 2) unfed and partially engorged tick tissues (salivary gland, midgut, ovary) from PA only with the DNeasy Blood and Tissue kit catalog # 69506 (Qiagen, Valencia, CA) using th standard protocol provided by the manufacturer. Before processing, the ticks were sterilized by two rounds of subsequent washing in 10% bleach, 70% ethanol, and sterile phosphate-buffered saline (PBS) (Biosciences, Cat#R028, St. Louis, MO USA). DNA concentration and purity were analyzed using Nanodrop, and the extracted DNA samples were stored at −20 °C until further use.

### 4.3 Illumina library preparation and 16S rRNA sequencing

Illumina DNA library preparation and 16S sequencing were performed by MR DNA, Shallowater, TX, USA. V1-V3 variable region of 16S rRNA genes was amplified using the forward primer 27F (5’-GAGTTTGATCNTGGCTCAG-3’) and the reverse 519R (5’- GTNTTACNGCGGCKGCTG-3’) with a barcode on the forward primer. The HotStarTaq Plus Master Mix Kit (Qiagen, USA) was used with following PCR conditions - 94°C for 3 minutes, followed by 30 cycles of 94°C for 30 seconds, 53°C for 40 seconds and 72°C for 1 minute, after which a final elongation step at 72°C for 5 minutes was performed. Amplified products were checked on 2% agarose gel to confirm the appropriate size and intensity of bands. On the basis of molecular weight and DNA concentrations, equal proportions of multiple samples were pulled together and purified by using calibrated Ampure XP beads. Purified PCR products were used to prepare Illumina DNA library. The quality of the DNA libraries was confirmed by lab-on-chip analysis using the Bioanalyzer (Agilent Technologies, Inc. Santa Clara, CA, USA), and then 16S sequencing was performed on Illumina MiSeq platform at MR DNA, Shallowater, TX, USA. Three biological replicates of each of the controls were used. Controls used were DNA extraction blank control, negative control (buffer), negative control (sterile water), No template control, Positive DNA extraction control [62-63] (commercially available Mock Microbial Community Standard, ZymoBIOMICS catalog # D6306).

### 4.4 Data Processing

Quantitative Insights into Microbial Ecology (QIIME 2, https://qiime2.org) was used for sequence analysis. Raw fastq files were processed by fastq processor available on the MR DNA website, which provided the files compatible with the Earth microbiome project (EMP) paired-end format. Then “Atacama soil microbiome” tutorial (website link - : https://docs.qiime2.org/2021.2/tutorials/atacama-soils/) and “moving pictures” tutorial (website link -: https://docs.qiime2.org/2021.2/tutorials/moving-pictures/) were followed to process the sequencing data. A total of 17,258 and 36,358 raw reads were obtained before denoising step. DADA2 [64] was used for trimming, primer sequence removal, sequence denoising, paired-end merging, filtering of chimeric sequences, singleton removal, and sequence dereplication. This step yielded 5,000 sequences from each tissue sample, while 16,000 sequences from each of the whole tick samples for rarefaction curves. The rarefaction curve is getting leveled out, suggesting that collecting additional sequences beyond that sampling depth would not observe additional reads. Minimum overlap of 50 bases was used for paired end merging. Resultant sequences set obtained after DADA2 processing were aligned by MAAFT (ver.7) [65], and then a phylogenetic tree was created by using FastTree (ver. 2.1) [66]. Greengenes 13_8 99% OTU database [67] was used to train the Naïve Bayes classifier, to which the represented sequences were compared and a 97% sequence similarity was put as a cut off for taxonomic classification. Network correlation maps were inferred based on the Sparse Correlations for Compositional data (SparCC) approach [68]. This approach uses the log-transformed values to carry out multiple iterations, which subsequently identifies taxa outliers to the correlation parameters [69]. Raw sequences were submitted to the NCBI read under SRA database and obtained accession number # PRJNA663181.

### 4.5 Statistical Analysis

To measure α-diversity, different indices such as Faith’s phylogenetic diversity (faith_pd) and Pielou’s community evenness (pielou_e) were used. Faith’s phylogenetic diversity (faith_pd) is an unweighted measure of phylogenetic distance of observed sequences; and Pielou’s community evenness (pielou_e) measures how evenly bacterial species are distributed within a community. Kruskal-Wallis non-parametric tests (p ≤0.05) were performed to determine statistical significance of alpha-diversity metrics by using QIIME 2. Weighted and unweighted UniFrac Metrics [70] were used for β-diversity analysis. EMPeror [71] was used for visualization of principal coordinate analysis (PCoA) plot, and PERMANOVA tests (p ≤0.05) were used to test the statistical significance of β -diversity measurements.

## 5. Conclusions

This study has shown significant differences in the microbiome of *Ix. scapularis* ticks collected from Northeastern (New York and Pennsylvania) and Southern (Oklahoma and Louisiana) states. These results provide an insight into the microbiome of NY, PA, OK, LA tick populations and interplay between pathogenic and endosymbiotic rickettsiae. The question remains: what are the drivers behind these variations among the microbiome composition and diversityã This question warrants further investigation such as why Oklahoma ticks contain comparatively much higher level of Firmicutes while ticks from all other locations included in this study contain almost negligible Firmicutes. In further analysis at the species level, it was revealed that these Firmicutes from Oklahoma ticks possess bacterial species such as *Jeotgalicoccus* sp. and *Staphylococcus lentus*. Do these species restrict *B. burgdorferi* s.l. in Oklahoma ticksã Similarly, the level of Bacteroidetes is much higher in Northeastern ticks to almost insignificant in Southern ticks. Further analysis revealed that these Bacteroidetes possess bacterial species such as *Sphingobacterium faecium, Flavobacterium* sp., *Sphingobacterium* sp., *Flexibacter* sp. and *Flavobacter* spp. Do these Bacteroidetes interact with spirochetes to colonize Lyme disease causative agent (i.e. *B. burgdorferi* s.l.) into Northeastern ticksã Probable competitive interaction between *R. buchneri* and *B. burgdorferi* s.l. have surfaced in this study subject to be further investigated by dysbiosis experiments and bigger sample size.

## Author Contributions

Conceived and designed the experiments: DK, LPD, and SK. Performed the experiments: DK, LPD and SK. Analyzed the data: DK, AA and SK. Contributed reagents/materials/analysis tools: DK, EM, KO, RSO, ME, and SK. Wrote the paper: DK and SK. All authors read and approved the final manuscript.

## Institutional Review Board Statement

All animal experiments were performed in strict accordance with the recommendations in the Guide for the Care and Use of Laboratory Animals of the National Institutes of Health. The protocol for the blood-feeding of field-collected ticks was approved by the Institutional Animal Care and Use Committee of the University of Southern Mississippi (protocol # 15101501.1).

## Informed Consent Statement

All authors read and approved the manuscript for publication.

## Data Availability Statement

Data supporting the conclusions of this article are included within the article and its additional files. The raw datasets used and analyzed for the present study are available from the corresponding author upon reasonable request.

## Acknowledgments

We thank Abdulsalam Adegoke and Raima Sen for their technical assistance in extraction of genomic DNA.

