## Supplementary material for "An exploratory study on the microbiome of northern and southern populations of *Ixodes scapularis* ticks predicts changes and unique bacterial interactions": Figure S1

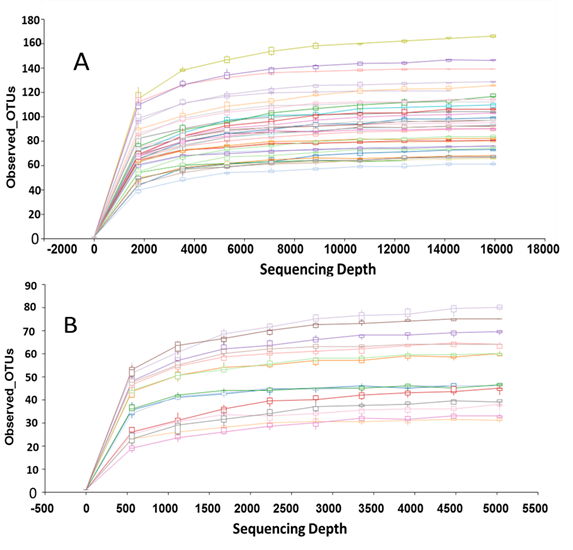
Figure S1: Rarefaction Curve. A) for whole ticks collected from Louisiana (LA), New York (NY), Pennsylvania (PA) and Oklahoma (OK). Data from sequence reads were rarefied to 16000 sequences. B) For unfed and partially fed ticks tissues (salivary gland, midgut, ovary). These ticks were field collected ticks from Pennsylvania. Unfed ovary data is not available. Data from sequence reads were rarefied to 5000 sequences. All samples contain at least three biological replicates.
